# wwLearning the language of proteins and predicting the impact of mutations

**DOI:** 10.1101/2024.04.05.588133

**Authors:** Bin Hu, Michal Babinski, Kaetlyn Gibson, Po-E Li, Valerie Li, Patrick Chain

## Abstract

Mutations in proteins directly impact their structure and function. Understanding the “language” of proteins, or the sequence to function (genotype-phenotype) relationship has many real-world applications. One set of applications includes those in biodefense, such as biological threat detection and biosurveillance, antibody engineering, and medical countermeasure development. In this study, we present a novel language model-based approach that can rapidly analyze vast collections of sequences, and make near real-time functional predictions that compare favorably to those made using conventional bioinformatic and experimental methods. Our findings reveal that tailored protein language models can predict protein mutation phenotypes, such as binding affinity or level of expression, when they are trained with high-throughput functional data. Protein language models applied to viral genomes can also discern the lineage within a family (e.g., sarbecovirus sequences). Coupled with sequenced-based biosurveillance, this type of model may provide early warning signals of potential zoonotic spillovers (i.e. host jumping) or “escape” from existing medical countermeasures posed by novel mutations. This research not only underscores the potential of ML and language models in addressing pressing challenges in understanding the mapping of sequence to function, but further elucidates their potential application in accelerating the response to biological threats as they evolve.

## Introduction

Proteins are fundamental to nearly all biological functions, playing crucial roles in cellular, pathological, and immunological processes. Those functions are achieved by proteins that are defined by amino acid sequences and configured into specific three dimensional structures. When under selection, protein mutations that lead to gain or loss of function, can be selected for and can result in cancer, immune escape, or loss of binding and ineffectiveness of medical countermeasures. Detection of genetic mutations can be readily performed using genomic sequencing, however linking observed mutations to phenotypic impact typically requires additional biochemical characterization – a process that can take months to experimentally discern. To cost-effectively characterize the impact of many different mutations, high throughput assays are required, together with the sequences of the mutants. For example, deep mutation scanning (DMS) (Fowler and Fields, 2014) and reprogrammed yeast mating (Younger et al., 2017) can generate and characterize tens of thousands of mutated protein sequences in a few months.

High throughput experiments have been applied to study SARS-CoV-2, a member of the betacorona virus family and the causative pathogen of COVID-19. The infection of SARS-CoV- 2 is initiated with the binding of the Spike protein of the virus through its receptor binding domain (RBD) to the human ACE2 receptor (Lan et al., 2020), which makes RBD the primary target of COVID-19 therapeutic antibodies and vaccines (Kleanthous et al., 2021; Min and Sun, 2021; Yang et al., 2020). Mutations in the RBD can lead to changes in the virus pathogenicity as well as to evasion of medical countermeasures (reviewed in (Carabelli et al., 2023)). Some mutations change the antigenicity and other biochemical properties of RBD, which can lead to escape from antibody or small molecule therapeutics and result in lower vaccine efficacy, as demonstrated during the pandemic by the Omicron (B1.1.529) lineage of SARS-CoV-2 that escapes antibody neutralization (Willett et al., 2022). In some rare events, gain of function mutations may allow a virus to switch its natural host and can lead to severe threat to public health.

With sufficient functional data paired with sequencing information, machine learning (ML) methods have been shown to be able to extract and interpret patterns found in biological sequences (i.e. language of proteins) and predict functions (reviewed in (Asnicar et al., 2023), see also (LeCun et al., 2015)). A popular ML method for sequence analysis is the long short term memory (LSTM) network originally used in natural language processing (NLP) and often used in language translation (Jurtz et al., 2017). LSTM has been applied to predict protein localization and secondary structure (Heffernan et al., 2017). Hie et al. demonstrated language models can be used to explore both the fitness and functional similarity of protein evolution (Hie et al., 2021).

More recently, LSTM has become gradually replaced by large language models (LLMs) that leverage transformers (Devlin et al., 2019). The training of LLMs consists of two phases: pre-training and fine-tuning. During pre-training, a model processes a massive text corpus with self-supervised learning tasks, usually predicting masked words in a given context, to learn the rich language representations, including vocabulary, grammar and semantics. The model is then fine-tuned with supervised learning tasks, such as sentiment classification, to improve performance. One benefit of the two phases of learning is that the pre-training can produce reusable foundation models that can be transferred to address new related problems, especially when these problems share similarities in grammar and vocabulary. Such advantages have attracted researchers to develop ProteinBERT (Brandes et al., 2022), TAPE (Rao et al., 2019), and other tools aimed at discovering complex patterns in protein data. More sophisticated machine learning models trained by many diverse sequences from across the tree of life (Doolittle, 1999), including UniRef sequences selected from UniProt and UniParc (Suzek et al., 2015), can predict protein structure with great accuracy (Akdel et al., 2022; Jumper et al., 2021; Lin et al., 2023; Rives et al., 2021). In one such example, researchers showed that biochemical properties of amino acids, secondary structure, and remote homology are captured through a transformer model (Rives et al., 2021). Rives et al. released the numerical representations of the amino acids, hereafter referred to as ESM embeddings (Rives et al., 2021).

We are interested in developing language models that can take the pathogen protein sequence as the input in order to predict biosecurity and public health related phenotypes, such as changes in pathogenicity, vaccine or therapeutic escape, and spill over potentials. Here we compare different model architectures to study the viral Spike RBD and the Spike-binding antibody sequences. We show that: 1) Protein language models can capture hidden patterns in protein sequences that relate to virus lineage and the host they are targeting; 2) Regression layers, which learn a mapping function from input features to a numerical output by adjusting parameters on training data, can be added to language models to predict biochemical properties of mutated proteins at accuracies comparable to wet bench experiments; 3) Protein language models can be a tool utilized during genomic surveillance efforts to help predict spillover events and escapes from therapeutics.

## Results

### Data collection

We collected and curated four datasets. The first, **outbreak dataset** containing 347,624 unique curated RBD sequences was obtained from both GISAID (Elbe and Buckland-Merrett, 2017) and the sequence reads archive (SRA) on February 3rd, 2023, with all duplicated sequences and sequences with ambiguous amino acids removed. This comprehensive key viral protein dataset represents sequences of circulating viruses under real-time evolution and selection during the pandemic. All major lineages of the virus are represented (Omicron: 52.70%, Delta: 35.71%, Alpha: 7.20%, **Figure 1A**). All 20 canonical amino acids are represented with Methionine (M) as an outlier, accounting for only 0.005% of total RBD amino acids and appearing in less than 1% of all the RBD sequences (**Figure 1B**). The second, **DMS dataset**, consists of 116,257 unique RBD sequences with measured dissociation constant (*K*_d_) to monomeric human ACE2 and 105,525 unique RBD sequences with expression levels measured from *in vitro* experiments (Starr et al., 2020). The third, **betaCov dataset**, includes 75 RBD sequences from SARS-CoV-2, including the original Wuhan-HU-1 strain (Wu et al., 2020), SARS-CoV-1 (Skowronski et al., 2005), middle east respiratory syndrome coronavirus (MERS-CoV, reviewed in (de Wit et al., 2016), and bat coronaviruses used in a recent RBD evolution study (Starr et al., 2022). The betaCov dataset represents RBD sequences that target a spectrum of mammals, including bat, pangolin, civet, camel, and human (**Table 1**). Some of the SARS-CoV-1 and MERS-CoV RBD sequences have been shown to bind to both human and camel receptors. We wished to evaluate whether the same type of machine learning approach used for predicting viral phenotype from RBD/spike protein sequence could also be used to predict antibody affinities to their target (in this case antibodies to the Spike protein). We collected a fourth **Ab dataset** 87,808 unique single chain variable fragment SARS-CoV-2 antibodies (scFV) sequences with measured binding affinity to the wild type SARS-CoV-2 RBD sequences generated in a yeast expression system (Engelhart et al., 2022). We split each of the above four datasets into training (80%) and testing (20%) sets with only the training dataset being used in the training process.

**Figure 1.**
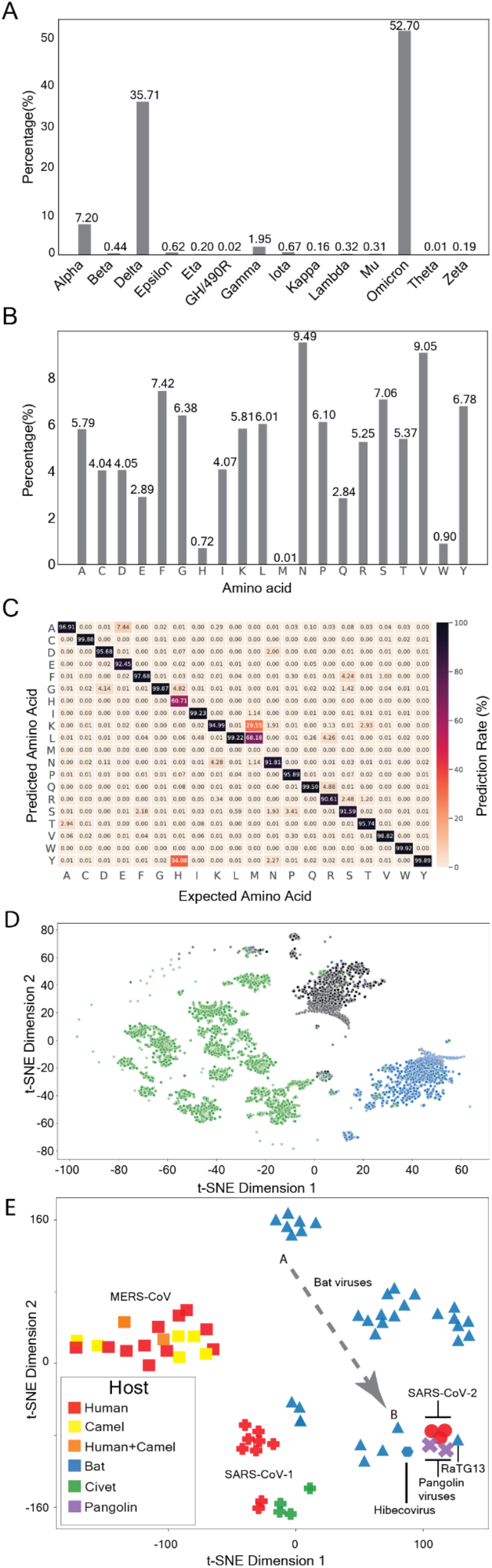
Outbreak dataset from real world SARS-CoV-2 sequences and language model result. **A.** Distribution of lineages covered in the outbreak dataset. **B.** Amino acid distribution of the RBD sequences. **C.** Language model with 320 dimensions for amino acid embedding and 10 attention heads after 100 epochs. X-axis: amino acids that are masked. Y-axis: amino acids predicted by the language model. Numbers in each row represent the prediction rate for the amino acid of that row. **D**. t-SNE plot of the BLSTM latent representation of three major SARS-CoV-2 lineages in the outbreak dataset with the ESM embedding. Black: Alpha, blue: Delta, green: Omicron. Omicron and Delta sequences are randomly down sampled to the same number of Alpha lineage sequences. **E.** Latent representations of virus sequence can serve as a radar for detecting potential public health threatening mutations. Squares: MERS-CoV, crosses: SARS-CoV-1, triangles: bat coronavirus, circles: SARS-CoV-2, X: pangolin coronavirus, hexagon: hibecorvirus. Red: isolated from humans, yellow: isolated from camels, orange: both human and camel, blue: isolated from bats, purple: isolated from pangolins, green: isolated from civets. A and B and the connecting dashed arrow t indicate a hypothetical mutation path that can lead to increased possibility to infect humans.

**Table 1.** Betacoronavirus used for clustering.

| Sequence Name | Accession | Variant | Host |
| --- | --- | --- | --- |
| RaTG13 | MN996532 | Bat Virus | Bat |
| ZXC21 | MG772934 | Bat Virus | Bat |
| ZC45 | MG772933 | Bat Virus | Bat |
| WIV16 | KT444582 | Bat Virus | Bat |
| Rs4231 | KY417146 | Bat Virus | Bat |
| WIV1 | KF367457 | Bat Virus | Bat |
| Rs7327 | KY417151 | Bat Virus | Bat |
| Rs4084 | KY417144 | Bat Virus | Bat |
| RsSHC014 | KC881005 | Bat Virus | Bat |
| LYRa11 | KF569996 | Bat Virus | Bat |
| As6526 | KY417142 | Bat Virus | Bat |
| Rs4237 | KY417147 | Bat Virus | Bat |
| Rs4247 | KY417148 | Bat Virus | Bat |
| Rs4081 | KY417143 | Bat Virus | Bat |
| 279-2005 | DQ648857 | Bat Virus | Bat |
| Rp3 | DQ071615 | Bat Virus | Bat |
| HKU3-8 | GQ153543 | Bat Virus | Bat |
| HKU3-13 | GQ153548 | Bat Virus | Bat |
| HKU3-1 | DQ022305 | Bat Virus | Bat |
| Longquan-140 | KF294457 | Bat Virus | Bat |
| GX2013 | KJ473815 | Bat Virus | Bat |
| Yunnan2011 | JX993988 | Bat Virus | Bat |
| Shaanxi2011 | JX993987 | Bat Virus | Bat |
| HuB2013 | KJ473814 | Bat Virus | Bat |
| YN2013 | KJ473816 | Bat Virus | Bat |
| JL2012 | KJ473811 | Bat Virus | Bat |
| Rf1 | DQ412042 | Bat Virus | Bat |
| 273-2005 | DQ648856 | Bat Virus | Bat |
| HeB2013 | KJ473812 | Bat Virus | Bat |
| Rf4092 | KY417145 | Bat Virus | Bat |
| RmYN02 | EPI_ISL_412977 | Bat Virus | Bat |
| BM48-31 | NC014470 | Bat Virus | Bat |
| BtKY72 | KY352407 | Bat Virus | Bat |
| Hp-BCoV_Zhejiang_2013 | YP_009072440.1 | Hibecovirus | Bat |
| SARS-CoV-1_SZ3_PC03 | AY304486 | Sars-Cov-1 | Civet |
| SARS-CoV-1_SZ13_PC03 | AY304487 | Sars-Cov-1 | Civet |
| SARS-CoV-1_SZ1_PC03 | AY304489 | Sars-Cov-1 | Civet |
| SARS-CoV-1_PC4-127_PC04 | AY613951 | Sars-Cov-1 | Civet |
| SARS-CoV-1_PC4-13_PC04 | AY613948 | Sars-Cov-1 | Civet |
| SARS-CoV-1_PC4-137_PC04 | AY627045 | Sars-Cov-1 | Civet |
| SARS-CoV-1_Urbani_HP03L | AY278741 | Sars-Cov-1 | Human |
| SARS-CoV-1_GD01_HP03L | AY278489 | Sars-Cov-1 | Human |
| SARS-CoV-1_BJ02_HP03M | AY278487 | Sars-Cov-1 | Human |
| SARS-CoV-1_GZ-C_HP03L | AY394979 | Sars-Cov-1 | Human |
| SARS-CoV-1_HGZ8L1-A_HP03E | AY394981 | Sars-Cov-1 | Human |
| SARS-CoV-1_Sino1-11_HP03L | AY485277 | Sars-Cov-1 | Human |
| SARS-CoV-1_GD03T0013_HP04 | AY525636 | Sars-Cov-1 | Human |
| SARS-CoV-1_Sin852_HP03L | AY559082 | Sars-Cov-1 | Human |
| SARS-CoV-1_GZ0402_HP04 | AY613947 | Sars-Cov-1 | Human |
| SARS-CoV-2 | MN908947 | Sars-Cov-2 | Human |
| SARS-CoV-2-related | QRN78347 | Sars-Cov-2 | Human |
| SARS-CoV-2-related | QWK65230 | Sars-Cov-2 | Human |
| Pangolin_GD-consensus | EPI_ISL_410544 | Pangolin Virus | Pangolin |
| Pangolin_GX-P2V | EPI_ISL_410542 | Pangolin Virus | Pangolin |
| MERS-CoV-related | ALA49363.1,<br>ATG84833.1 | MERS-CoV | Camel,<br>Human |
| MERS-CoV-related | ALA49374.1,<br>AHZ58501.1 | MERS-CoV | Camel,<br>Human |
| MERS-CoV-related | ALA49836.1,<br>ANF29239.1 | MERS-CoV | Camel,<br>Human |
| MERS-CoV-related | AKO69634.1 | MERS-CoV | Camel |
| MERS-CoV-related | ALA50067.1 | MERS-CoV | Camel |
| MERS-CoV-related | AHY22545.1 | MERS-CoV | Camel |
| MERS-CoV-related | ASU90329.1 | MERS-CoV | Camel |
| MERS-CoV-related | ASU90527.1 | MERS-CoV | Camel |
| MERS-CoV-related | AID55088.1 | MERS-CoV | Human |
| MERS-CoV-related | ANC28634.1 | MERS-CoV | Human |
| MERS-CoV-related | ALK80291.1 | MERS-CoV | Human |
| MERS-CoV-related | AGV08379.1 | MERS-CoV | Human |
| MERS-CoV-related | ALJ54518.1 | MERS-CoV | Human |
| MERS-CoV-related | ALJ54491.1 | MERS-CoV | Human |
| MERS-CoV-related | AKN24803.1 | MERS-CoV | Human |
| MERS-CoV-related | AKL59401.1 | MERS-CoV | Human |
| MERS-CoV-related | ASY99842.1 | MERS-CoV | Human |
| MERS-CoV-related | ALJ54446.1 | MERS-CoV | Human |

### Learning from RBD sequences using a protein language model

Protein sequences are first converted to numerical representations (also known as embeddings) before they are processed by machine learning models. Several embedding methods exist, including k-mer cooccurrence-based methods such as GloVe (Pennington et al., 2014), a method using principal components of amino acid properties (Gelman et al., 2021), and the binary one-hot representation, where each amino acid is uniquely encoded as a vector with a single "hot" (1) position corresponding to its identity and all other positions set to "cold" (0). Considering its unique ability to accommodate partially overlapping biochemical properties in language models, we decided to learn the amino acid embeddings using a transformer-based language model – BERT (Devlin et al., 2019). We modified the original BERT model so that it learns amino acid embeddings with only one pre-training task of predicting the sequences of 15% randomly masked amino acids given the context of the RBD sequence. Using the outbreak dataset, the modified BERT model was able to predict the masked amino acid residues with an average correct prediction rate of 90% with 320 dimensions and 10 attention heads after 100 epochs of training (**Figure 1C**). That is, given the context of a sequence, this model can accurately predict the masked amino acid 90% of the time. This model underperforms in predicting methionine, which is not surprising given that methionine appears very rarely (0.005%) in the RBD sequences. The second poorest prediction is histidine, which is also the second lowest represented amino acid (0.72%) (**Figure 1B**). Training the modified BERT model also generated a new embedding that uses 320 numbers to represent an amino acid. Since this embedding is unique to the RBD sequences from the outbreak dataset, we call it RBD learned embedding.

### Predicting viral lineage and host from viral sequences

Since the lineage information of the outbreak dataset and the host information of the betaCov dataset are both available, we reasoned that the model can also be evaluated to see whether it can distinguish different RBD sequences by lineage and by host. We linked an ESM model to a bidirectional LSTM (BLSTM) model, hereafter called as ESM-BLSTM model. In this model, the ESM sequence embeddings are fed to a BLSTM model (Hochreiter and Schmidhuber, 1997). The last hidden layer of the BLSTM model, which captures the information extracted from an entire sequence, is decomposed to two-dimension using t-SNE (Maaten and Hinton, 2008) or UMAP (McInnes et al., 2020) for visualization. Since the SARS-CoV-2 Omicron, Delta, and Alpha lineages together account for 95.11% of the outbreak dataset used in this experiment (**Figure 1A**), we focused on these three lineages and down sampled Omicron and Delta sequences randomly so that three lineages are equally represented with 22,073 sequences each. **Figure 1D** plotted the decomposed final hidden state from the ESM-BLSTM model with the data labeled by lineage information, which accurately separates the sequences from different lineages without using any clustering method. Given the randomness in subsampling, we repeated the same experiment 12 times using t-SNE and UMAP, and the lineage separation is repeatedly found in all experiments (**Supplementary Figure 3**). The t-SNE and UMAP plots show clusters composed of sequences from different lineages or with unassigned lineages that appear to be more similar to sequences that were assigned to the same lineages. We checked the lineage composition and sample collection dates of three clusters mainly of Omicron lineage (**Supplementary Table 1**). We argue that the unknown lineages of sequences in a cluster may be assigned with the dominating lineage of that cluster. The collection dates of the three clusters showed different peaks, which may suggest different sublineages dominating the spreading of the virus of that time. However, additional data, including geographic information and local epidemiological data may be needed to interpret the peaks of collection dates associated with these clusters. We concluded that the ESM-BLSTM model can be used to assist sublineage assignment and identify sequences of high similarity regardless of the lineage assignment by other tools.

To evaluate whether the ESM-BLSTM model can also separate closely related viruses affecting different hosts, we investigated the last hidden layer of the model using the betaCov dataset. As shown in **Figure 1E**, betaCov sequences form several clusters with the bat coronaviruses more dispersed. Some bat coronaviruses are projected close to SARS-CoV-2, Pangolin coronavirus, and RaTG13, a close relative of SARS-CoV-2 (Zhou et al., 2020) which can be neutralized by convalescent sera from COVID-19 patients (Cantoni et al., 2022). In a hypothetical bat virus sequence surveillance effort, a trajectory from point A to point B over time may indicate a series of mutations gradually leads to increased risk for infecting humans (**Figure 1E**).

### Protein sequence to phenotype prediction

Given that our ESM–BLSTM model can accurately predict masked amino acids and extract pattern in the RBD sequences for lineage separation, we decided to adapt the ESM-BLSTM model to predict the binding affinities and expression levels in the DMS dataset (Starr et al., 2020). We compared the model performance in predicting quantitative results using different embedding methods and model architectures, including a fully connected neural network (FCNN), a graph convolutional neural network (GCN) based on the wildtype RBD 3D structure, BLSTM, ESM- or BERT-BLSTM (**Table 2**). We use the root of mean squared error (RMSE) as the performance indicator. RMSE indicates the average error from the entire testing run, thus a smaller RMSE indicates a better model performance. We found that ESM embeddings generally make more accurate predictions than using the RBD learned embeddings (**Table 2 and Figure 2**). FCNN alone (without BERT or BLSTM) was able to predict the binding affinities with RMSE=1.23 (**Figure 2A**). GCN performed slightly better (RMSE=1.18, **Figure 2B**). The best models in predicting binding affinity are 1) a BLSTM model with the last hidden status fed to the FCNN (**Figure 2C**); and 2) an ESM-BLSTM model (**Figure 2D**), where the initial ESM embeddings are updated throughout the training process along with the BLSTM component.

**Figure 2.**
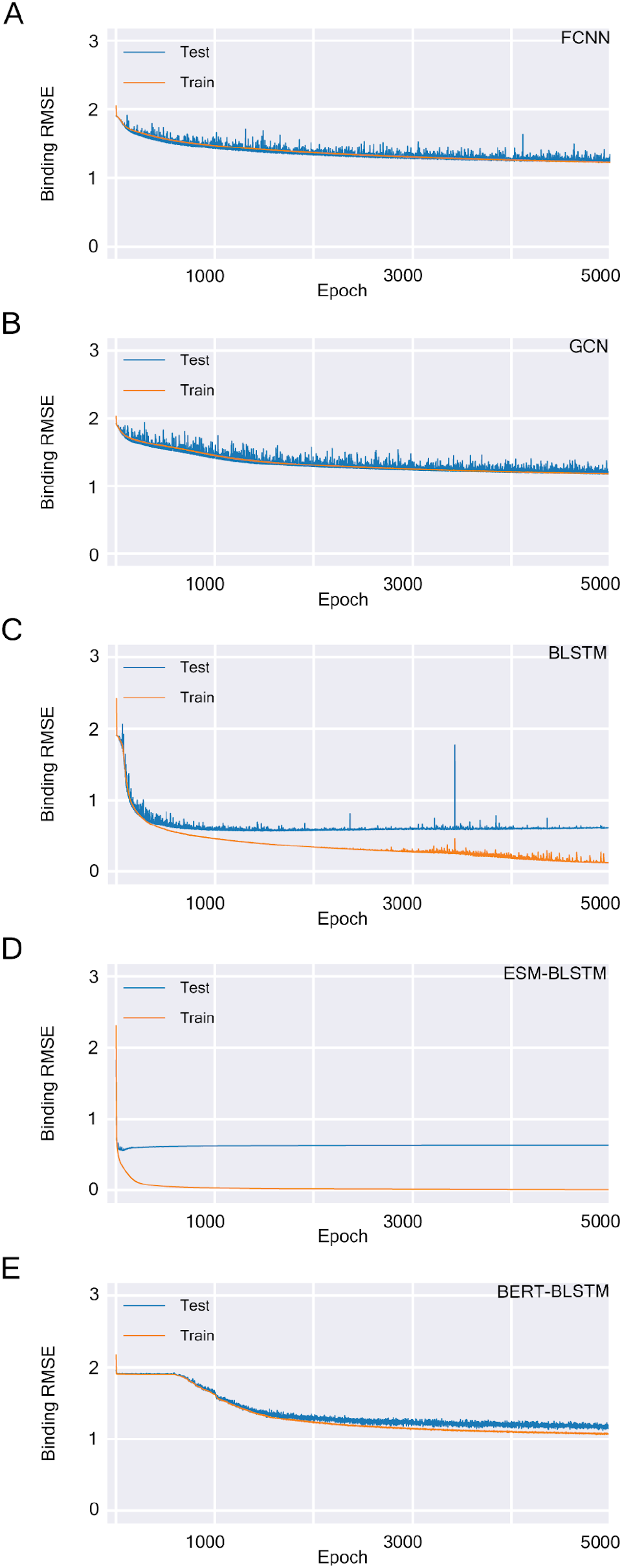
Predicting RBD-ACE binding affinities using different models with ESM embedding. **A.** Fully connected neural network. **B.** Graph convolutional network. **C.** Standalone BLSTM model. **D.** ESM-BLSTM mode with ESM embeddings updated through the training process. **E.** BERT-BLSTM model with ESM embeddings updated through the training process.

**Table 2.** Binding and expression prediction comparisons (architecture and embedding)

| Model Architecture | Embedding Method | Embedding Updated | Binding Error (RMSE) | Expression Error (RMSE) |
| --- | --- | --- | --- | --- |
| FCNN | ESM | No | 1.226022382705891 | 0.7325193013771152 |
| FCNN | RBD Learned | No | 1.9174587664223277 | 1.0517566963652063 |
| GCN | ESM | No | 1.1772057508817406 | 0.7319495048945327 |
| GCN | RBD Learned | No | 1.9066781674836124 | 1.0415343171336675 |
| BLSTM | ESM | No | 0.5639194085741418 | 0.4647852894674934 |
| BLSTM | RBD Learned | No | 1.912067333676975 | 1.0416382379402376 |
| ESM-BLSTM | ESM | Yes | 0.5565162630343344 | 0.4569100085850099 |
| BERT-BLSTM | ESM | Yes | 1.1222631753194006 | 0.7747551330511154 |
| BERT-BLSTM | RBD Learned | Yes | 1.8678186524258549 | 1.0392111455289788 |

Both models reach RMSE=0.56 during the training period. Similarly, we tested a BERT-BLSTM model, where the BERT model had the initial embeddings from ESM and allowed embedding updates during the training process. However, the BERT-BLSTM model did not generate a satisfactory prediction (RMSE=1.12 when initialized with ESM embedding, RMSE=1.87 when initialized with the RBD learned embedding, Figure **2E**). In another experiment to predict the *in vitro* RBD expression levels, the most accurate predictions were also found in BLSTM models with ESM embeddings and the ESM-BLSTM model (both at RMSE=0.46, **Table 2 and S Figure 5**).

### Antibody affinity prediction

While the use of the modified BERT model and the ESM-BLSTM model were found to successfully predict masked amino acids, separate virus lineages, and shows potential application in biosurveillance based on viral protein sequence data, we explored if the use of similar models could be extended to predict antibody binding phenotypes based on antibody sequences. We used the modified BERT language model to learn the embeddings of the Ab dataset and predict masked sequence information. However, the average prediction rate is only at 29.91% (**Figure 3A**). This significantly poor predictive performance is potentially due to the scFvs likely targeting different epitopes of RBD coupled with insufficient training data to learn patterns from all the different types of scFvs. Indeed, when we look at the last hidden state of the ESM- BLSTM model, we noticed clustering of antibodies, especially for those showing high binding affinities (**Figure 3C**). We used RMSE to evaluate the model performance and compared model RMSE with experimental RMSEs, which is calculated by taking the average of multiple measurements of the binding affinity of the same sequence as the true value. Applying an ESM- BLSTM model to predict the scFv binding affinities with the average RMSE 0.8, which is comparable to experimental RMSE (0.6). Considering the peak of the distribution of measured binding affinity is about 4 log_10_ nM (**Supplementary Figure 6**). The ESM-BLSTM model RMSE is 20% of a typical measurement, which is comparable to experimental error (15%). Taken together, the ESM-BLSTM model’s predictions are comparable to experimental results in accuracy.

**Figure 3.**
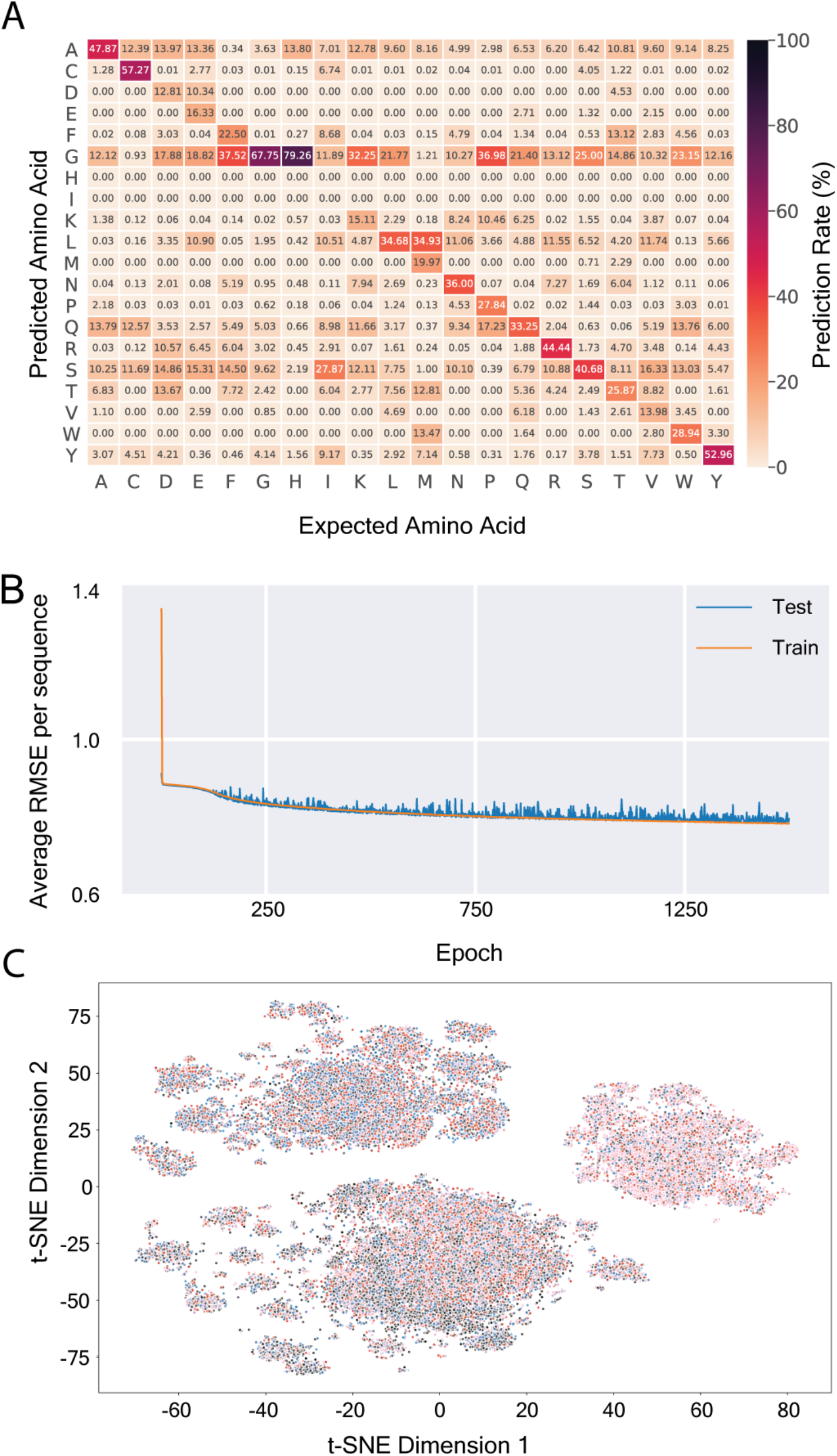
Antibody language model, binding affinity prediction, and clustering. **A.** Training results after 105 epochs. X-axis: amino acids that are masked. Y-axis: amino acids predicted by the language model. Numbers in each row represent the prediction rate for the amino acid of that row. **B.** Binding affinity predictions. **C.** Latent representation of the antibody sequences visualized in t-SNE. Black: 1st quantile, blue: 2nd quantile, red: 3rd quantile, pink: 4th quantile.

## Discussion

Genomic surveillance of SARS-CoV-2 during the pandemic has resulted in many millions of genomes sequenced. Such a rich collection from both real-world genome surveillance of SARS-CoV-2 and high throughput experiments have enabled bioinformatics and machine learning (ML) approaches to investigate the evolution and spread of SARS-CoV-2 genomes with successful examples including lineage assignment (O’Toole et al., 2021), identification of mutations of the SARS-CoV-2 associated with fitness (Obermeyer et al., 2021), and patterns of long COVID from electronic health records (Zhang et al., 2023). We trained a BERT protein language model using the outbreak dataset. This model was able to predict randomly masked amino acids in the RBD sequences with great accuracy. One potential application of such a model is to predict missing amino acids due to low coverage from sequencing and detect sequences with important mutations that significantly deviates from the model. We also found that ESM-BLSTM model can help differentiate lineages of SARS-CoV-2 and predict binding affinities (host target and antigen) and expression levels with accuracy comparable to high-throughput experiments. In addition, we showed that machine learning models can potentially be used for sequence based biosurveillance and early detection of mutations that may lead to host jumps.

A key challenge in studying proteins using machine learning is to embed protein sequences with distributed representation that allows capture of both known and unknown biochemical properties. As reviewed in (LeCun et al., 2015), deep-learning utilizing distributed representations allows exponential advantages over classic learning algorithms in both generalizations beyond the training dataset and potential for composing layers of representation. Compared to traditional embedding methods, such as one-hot (20 dimensions) and biochemical property based methods (e.g. AAindex-PCA method with 19 dimensions used by (Gelman et al., 2021)), LLM methods can capture additional information from the pre-training process, including shared properties as shown in (Rives et al., 2021). Our pre-training focused on predicting the masked amino acids given the context of RBD sequences from different lineages. When the BERT model was trained using only 24 dimensions, it performed poorly in predicting the masked amino acids (**Supplementary Figure 1**), which may be caused by the lower number of dimensions not capturing the nuances found in different amino acids. Increasing it to 320 dimensions (**Figure 1C**), same as in ESM embeddings, provides a much improved embedding. Further increasing the embedding to 768 embeddings, same as used in the BERT model for modeling human language, did not bring noticeable improvement in predicting the masked amino acids, although it doubled the computing time (**Supplementary Figure 2**). Although the pre-training achieved reasonably good accuracy, the performance of the affinity and expression level predictions were poor when the amino acid embedding learned from this process was used in downstream predictions. This contrast revealed the difference between the pre-training and fine tuning tasks as shown in (Peters et al., 2019). The greatly improved performance of the same fine tuning tasks using ESM embedding also suggests that the performance of the model largely depends on the amount of training data. However, since the training data used by ESM consists of nearly all known available sequences from the tree of life with uneven representations in different branches (Rives et al., 2021), which may cause inherent bias in the learned embeddings. How to quantify the inherent bias in embeddings remains an open question.

Our BERT model approach focuses on learning the language used by a specific protein of a virus and achieved impressive results in predicting masked amino acids. However, when used in biochemical property predictions, the thus learned amino acid embeddings underperforms and even showed over fitting (**Figure 4A**). The superior performance of the ESM-BLSTM model in predicting binding and expression levels (**Table 2**) demonstrates that ESM embedding— although may be biased by the uneven representation of different proteins in the tree of life— excels at capturing the biochemical properties of different amino acids. One obvious method to correct such bias is finetuning the ESM embedding through adding training tasks such as binding and expression predictions that can also update the initial embeddings. However, to our surprise no improvement in binding and expression predictions are found when ESM embeddings were updated during the training process (**Table 2**).

**Figure 4.**
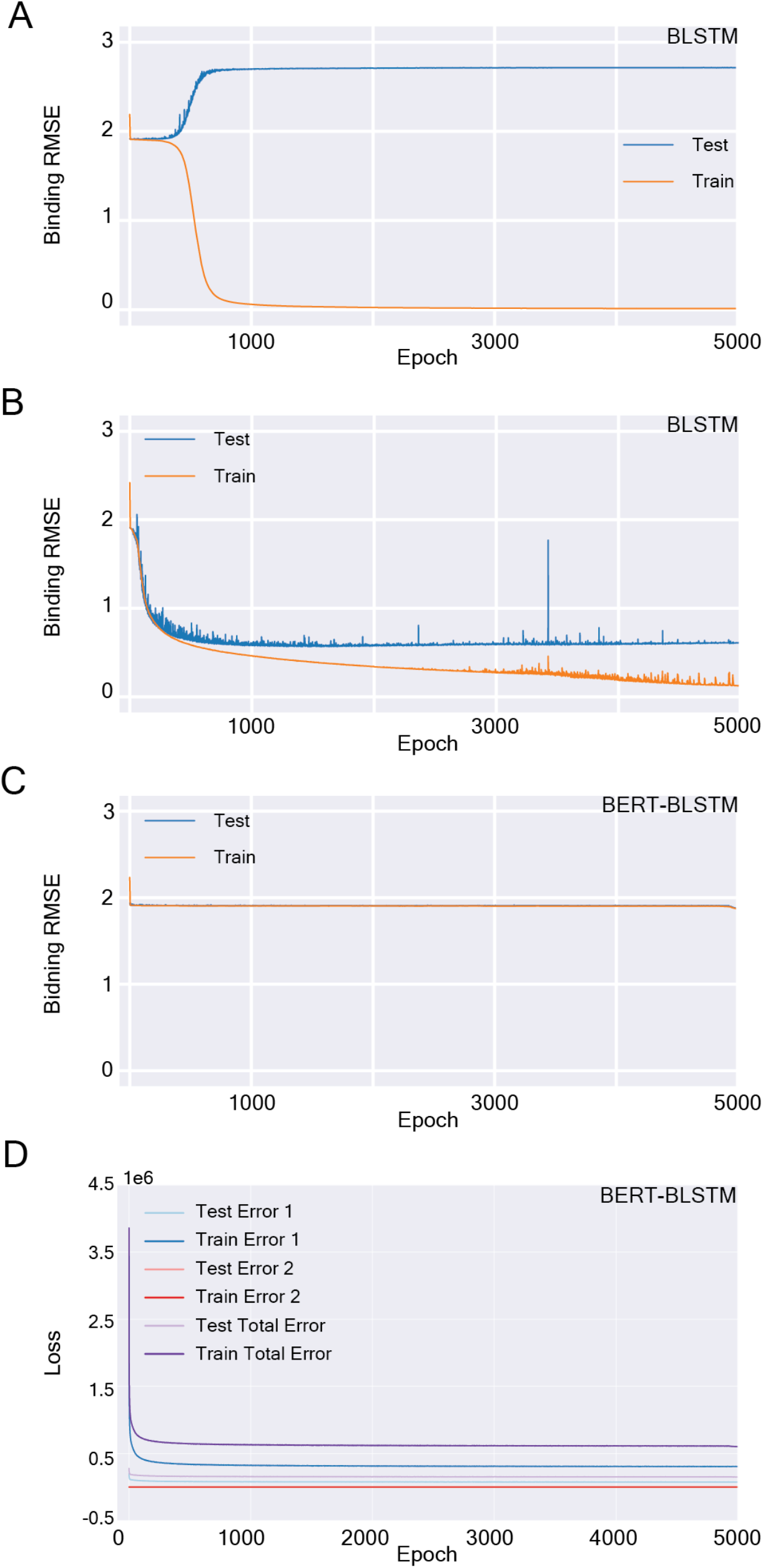
Model architecture and stability in predicting biochemical properties. **A.** RBD-ACE binding prediction using BERT-BLSTM model with fixed embeddings learned through the BERT masked language model. **B.** RBD-ACE binding prediction using ESM- BLSTM model with fixed ESM embeddings**. C.** RBD-ACE binding prediction using BERT- BLSTM model with embedding update with fine tuning. **D.** RBD-ACE binding affinity prediction using the BERT-BLSTM model with both pretraining (error 1, from the masked language model) and fine tuning error (error 2).

Switching to the ESM embedding using the same model, the RMSE improved to about 0.7 and showed over-fitting after ∼300 epochs (**Figure 4B**). However, the overfitting was much less than compared to the learned RBD embedding, which again suggests that the ESM embeddings are more accurate in capturing the hidden properties in different amino acids. Slightly better performance (RMSE ∼0.5) can be achieved if the BLSTM model is connected to the BERT model so that back propagation can update the learned RBD embeddings (**Figure 4C**). Interestingly, the BERT-BLSTM with updated embeddings didn’t show obvious overfitting until ∼1000 epochs. In this model, error from the masked language was not used to update the embeddings.

One possible method to further improve the model is to minimize both the pretraining masked language model error (*e*_1_) and the fine-tuning affinity prediction error (*e*_2_) simultaneously during the training process. To do that, we define the total error (*e_t_*) for each epoch as:

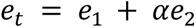

where α is weight coefficient for the two error terms and set to 1 since these two terms are in the same order. As shown in **Figure 4D**, this approach makes the model more stable.

The BERT-BLSTM model can predict mutated SARS-CoV-2 RBD binding to ACE and expression level and the anti-RBD antibody binding affinity at similar accuracy as the experiments. Compared to simpler method architecture, such as FCNN, GraphSAGE or BLSTM only method, more sophisticated architecture such as BERT-BLSTM, provides more stable predictions (**Figure 2**). The advantage of BERT-BLSTM over standalone BLSTM may come from the transformer architecture utilized in BERT, which relies heavily on multi-head attention networks (Vaswani et al., 2017) to offer enhanced proficiency in capturing context information over extended sequences. Our result shows great potential for generalized protein phenotype prediction using genotype information through a combination of LLM and high-throughput experimentation.

Graph convolutional network (GCN) based on protein structure abstraction (Gelman et al., 2021) provides a potential path towards the explainability of machine learning models. Through investigating the learned weights associated with each amino acid (vertex) and edges (distance from other amino acids) in GCN, one can trace back to the protein structure and identify key amino acids contributing to a specific biochemical function, such as receptor and immune molecule bindings. However, predictions from our GCN model are not better than the predictions from BERT-BLSTM or standalone BLSTM (**Figure 2**), which may be caused by the fixed graph structure used in our model even though the input RBD sequences are all different. To test this hypothesis, we plan to use ML-based protein folding tools such as AlphaFold-2 (Jumper et al., 2021) and ESMFold (Lin et al., 2023) to update the graph structures of the mutated RBDs in GCN.

The latent representation of the RBD sequences in three major lineages of SARS-CoV-2 (**Figure 1D**) provides satisfactory separation of these lineages and suggests ML may be applied in pathogen surveillance, especially zoonotic virus mutations. Our current model cannot separate different hosts, which may be caused by the host receptor protein variations not being included in the model. To address this limitation, a foundation model for studying biomolecule interactions is needed. Such a foundation model needs to incorporate both language models for sequence based feature and structure features so that a pair of biomolecules, such as a pathogen protein and a host receptor, can be studied simultaneously to better predict virus host selections and spillover events.

## Online Methods

### RBD sequence collection

All the SARS-CoV-2 spike protein sequences were downloaded from GISAID (Elbe and Buckland-Merrett, 2017) as of version 20230528 followed by MD5-based search for deduplication and removal of sequences with a length shorter than 600aa or containing any unknown residues (X), we obtained a dataset of 347,753 unique sequences. Using the reference sequence of RDB region (YP_009724390:319..541), we performed similarity search against the dataset using DIAMOND blastp algorithm (Buchfink et al., 2021) and extracted 347,624 RBD regions that align with the complete 223aa RBD region. A discrepancy of 347,409 records compared to 347,624 records exists in the EPI_SET due to 215 records missing from the GISAID COVID database. All the four datasets are available for download. Sequences and metadata from GISAID can be accessed using the EPI_SET ID: EPI_SET_240219yp.

### DMS data processing

All 195,081 initial entries for both the DMS binding and expression datasets were obtained from Starr et al., 2020. Entries with no binding or expression values were removed from the datasets. After selecting for nonsynonymous mutations, deduplication was applied by averaging the binding or expression value per amino acid substitution and deleting the duplicate entries. Amino acid substitutions were then applied to the reference sequence SARS-CoV-2 (MN908947) for each entry. The final number of entries with unique nonsynonymous sequences per dataset were 105,525 for binding and 116,257 for expression.

### Alpha-seq data processing

The raw Alpha-seq dataset was obtained from Engelhart et al., 2022. All entries missing a predictive affinity, values that estimate the protein-protein dissociation constant, were removed from the dataset. All sequences were then parsed through to find duplicates which were then placed in a set, and a mean affinity was given to each unique set. Then, only the first sequence in each set was kept with the corresponding mean affinity. The final number of unique sequences with mean affinities was 87,807.

### Beta-corona data processing

A majority of the sequences from the Beta-corona dataset were obtained from Starr et al., 2020. The MERS sequences were gathered from querying the accession number ALA49374.1 through BLASTp. After obtaining the full amino acid sequences, we manually extracted the RBD region and combined it with the sequences from Starr. The final number of sequences was 75.

### ML code

The original BERT model was designed with training focusing on understanding human languages (Devlin et al., 2019). To learn the protein language, we removed sentiment prediction and sentence tagging in BERT. The implementation of the BERT model is based on the source of the annotated Transformer(https://nlp.seas.harvard.edu/annotated-transformer/). The BERT model is used to process input sequences, with an extension Protein Language model that applies a simple linear model to map hidden features to vocabulary space, allowing for prediction of proteins. A custom protein vocabulary was used to tokenize the proteins passed to the BERT model. Attempts using more complex, and more computationally expensive versions of the modified BERT model did not yield noticeable benefit to the prediction accuracy. Tests were performed by increasing the embedding size from 320 to 768, and by increasing the attention heads from 10 to 12 (**Supplementary Figure 1**). To predict biochemical properties of the embedded sequences, the BLSTM architecture was applied to make regression predictions. The BLSTM is an extension of the LSTM model that processes data in both the forward and backwards direction. This allows the RBD sequence domain to be represented over a larger context window, where each element of the sequence is effectively considered. All the code used in this study is available in https://github.com/hubin-keio/Spike_NLP. In the GCN model, amino acids are represented as vertices, and edges between two vertices are assigned widths based on the Euclidean distance in a contact map built from the beta-carbons from the crystallography structure of the wild type SARS-CoV-2 RBD, similar to (Gelman et al., 2021). We used the GraphSAGE model implemented in the Deep Graph Library (Wang et al., 2020). To further explore DMS binding and expression prediction, the Graph Sample and Aggregation (GraphSAGE) model was used. The GraphSAGE model generates embeddings by sampling and aggregating features from nodes’ local neighborhood, making it unique from other neural networks that need the entire graph to generate embeddings. The ESM model (https://github.com/facebookresearch/esm) was used to embed input sequences. The embedded input sequences were passed as input to BLSTM and GraphSAGE models to predict DMS binding and expression. All model training and inference were executed on A100 GPUs (Nvidia, Santa Clara, CA).

### Data Availability

All genome sequences and associated metadata in the outbreak dataset are published in GISAID’s EpiCoV database with the GISAID identifier: EPI_SET_240219yp. It is composed of 347,409 individual genome sequences with collection dates range from 2019-06-25 to 2023-05- 17; Data were collected in 202 countries and territories. To view the contributors of each individual sequence with details such as accession number, Virus name, Collection date, Originating Lab and Submitting Lab and the list of Authors, visit doi:10.55876/gis8.240219yp The reference sequence for SARS-CoV-2 used in this study is hCoV-19/Wuhan/WIV04/2019 (WIV04), the official reference sequence employed by GISAID (EPI_ISL_402124, https://gisaid.org/WIV04)

### Saving Embeddings

To generate pickle embedding files of both DMS binding and expression datasets, data is processed using a specified embedding method using either the ESM model or the saved weights of the best performing RBD-trained BERT model. Each sequence in the dataset is represented as an entry containing the sequence identifier, measured binding or expression value, sequence, and the corresponding embedded sequence based on the last hidden state of the embedding method. The entries are then accumulated into a list and subsequently saved as a pickle file.

## Acknowledgment

We gratefully acknowledge all data contributors, i.e., the Authors and their Originating laboratories responsible for obtaining the specimens, and their Submitting laboratories for generating the genetic sequence and metadata and sharing via the GISAID Initiative, on which this research is based. The authors appreciate funding from CDC, Los Alamos National Laboratory (LDRD Director Initiated Research, 20240734DI, 20210767DI, 20200732ER). Part of this work was funded by the DOE Office of Science through the National Virtual Biotechnology Laboratory, a consortium of DOE national laboratories focused on response to COVID-19, with funding provided by the Coronavirus CARES Act. This project has been funded in part with Federal funds under Interagency Agreement No. 22FED2200087IPD (RRJJ) between the Centers for Disease Control and Prevention, Influenza Division and the US Department of Energy, Los Alamos National Laboratory. The GPU cluster was purchased by the bioassurance funding through DOD. This work is approved for public release under LA-UR-24-21857.

