## Supplementary Figures and Tables for "wwLearning the language of proteins and predicting the impact of mutations"

**Supplementary Material**

**S Figure 1. Language model training progression and different embedding size and heads (embedding size 24, 12 heads). A.** Amino acid prediction accuracy. **B.** Training loss in predicting masked amino acids. **C.** Training Accuracy in predicting masked amino acids.

**A**

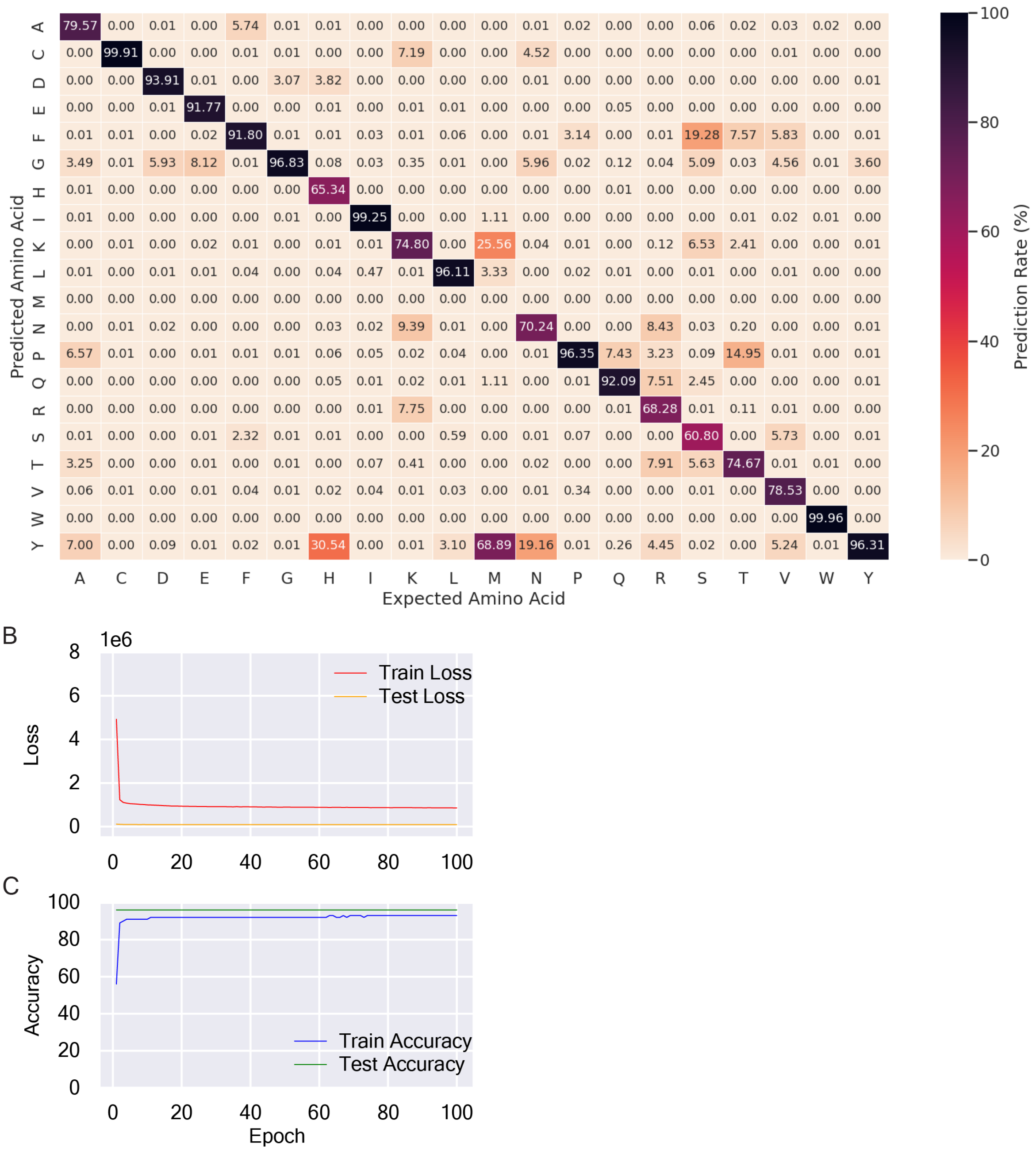

**S Figure 2. Language model training progression and different embedding size and heads. A.** Amino acid prediction accuracy (embedding size 768, 12 heads). **B.** Training loss in predicting masked amino acids (embedding size 768, 12 heads). **C.** Training accuracy in predicting masked amino acids (embedding size 768, 12 heads). **D.** Training loss in predicting masked amino acids (embedding size 320, 10 heads). **E.** Training accuracy in predicting masked amino acids (embedding size 320, 10 heads).

**A**

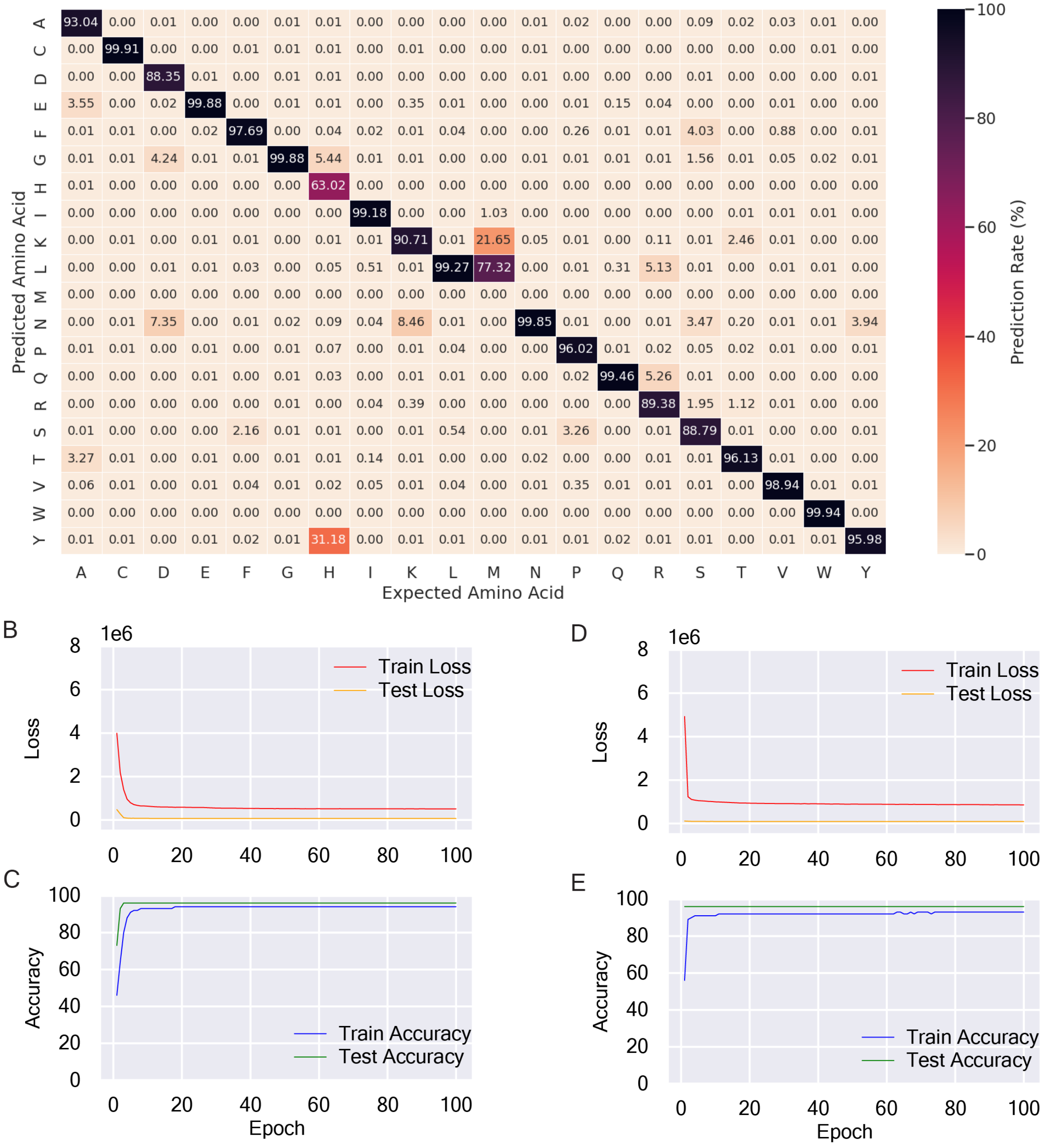

### S Figure 3. Additional t-SNE and UMAP visualization of lineage separation

A

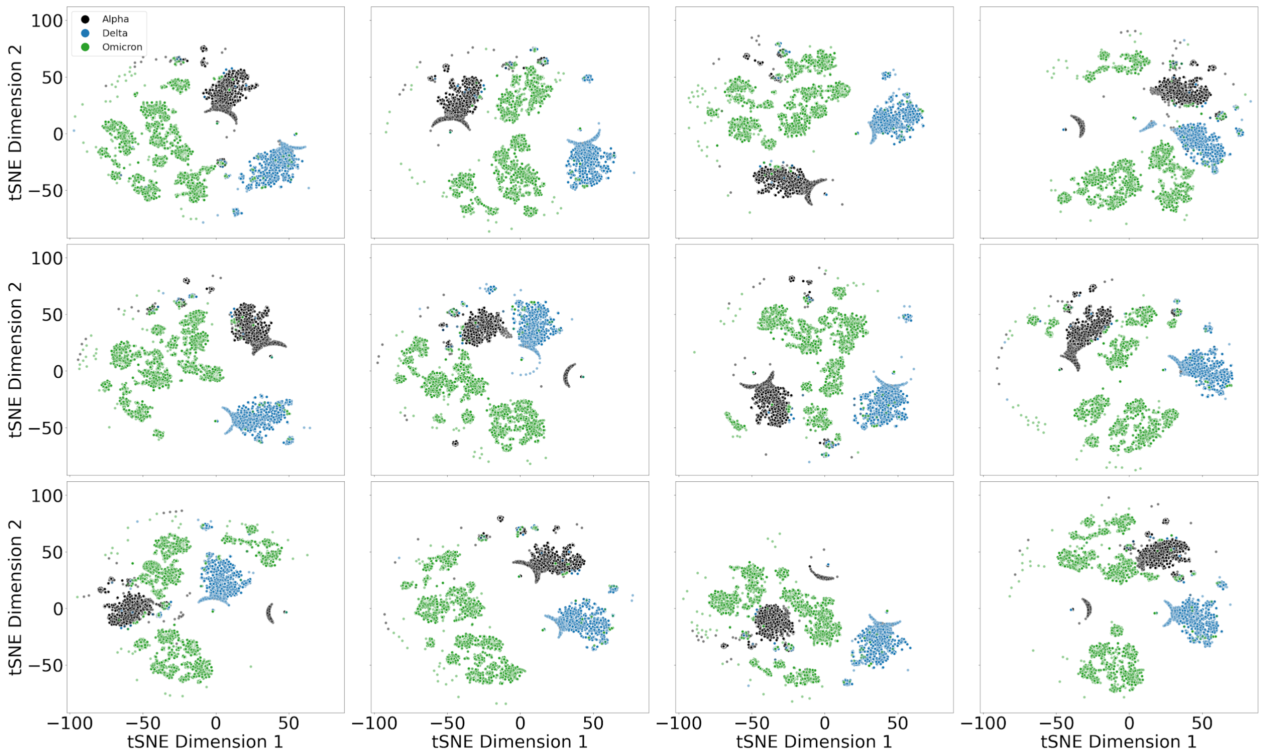


B

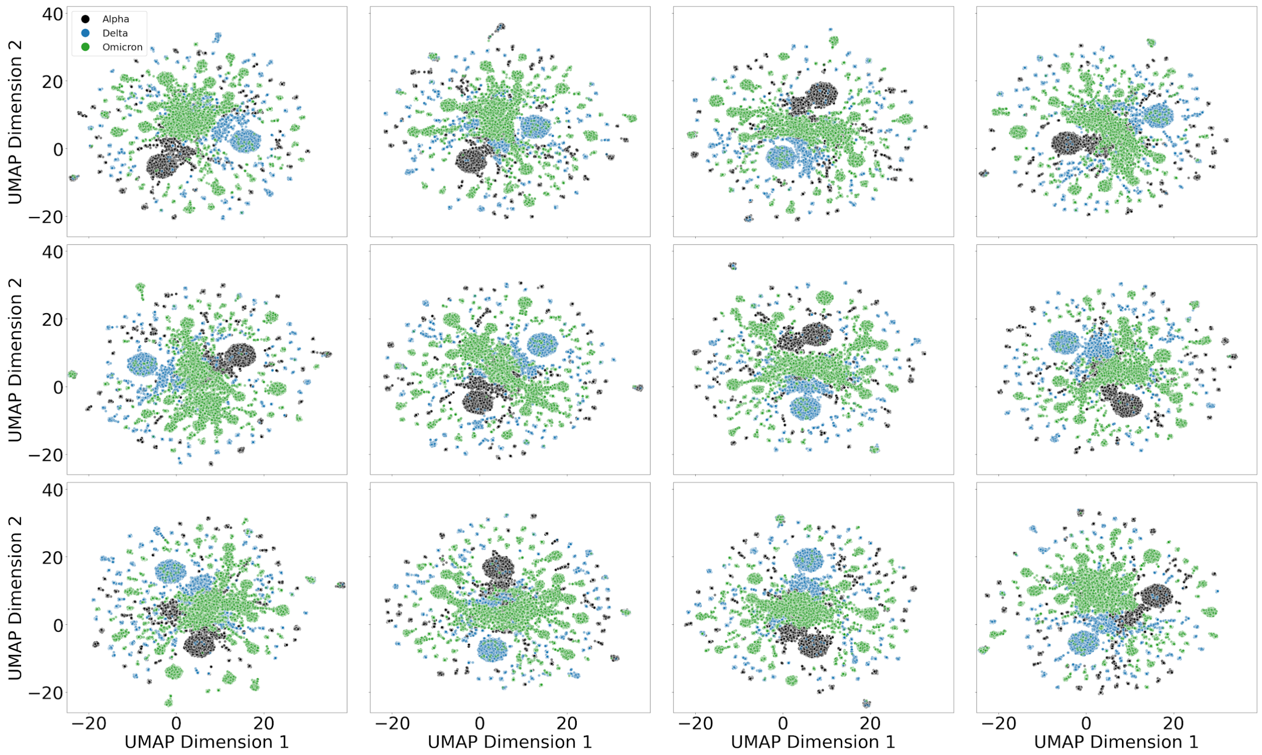

**S Figure 4. Mapping Collection Date to Omicron Clusters. A.** t-SNE plot with highlighted clusters. **B.** Collection date time plot for cluster 1. **C.** Collection date time plot for cluster 2. **D.** Collection date time plot for cluster 3.

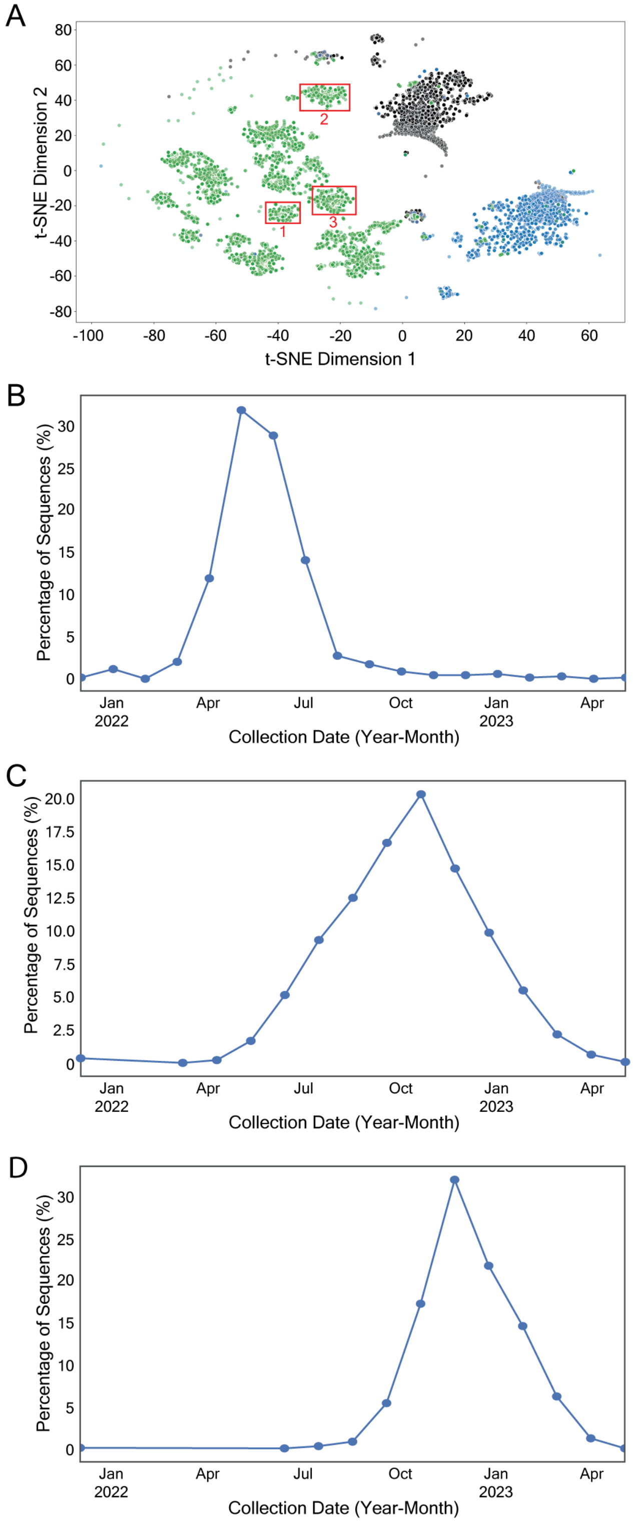

**S Figure 5. Predicting RBD expression levels using different models. A.** Fully connected neural network with ESM embedding. **B.** Graph convolution network with ESM embedding. **C.** Standalone BLSTM with ESM embedding. **D.** ESM-BLSTM model. **E.** BERT-BLSTM model (BERT connected to BLSTM).

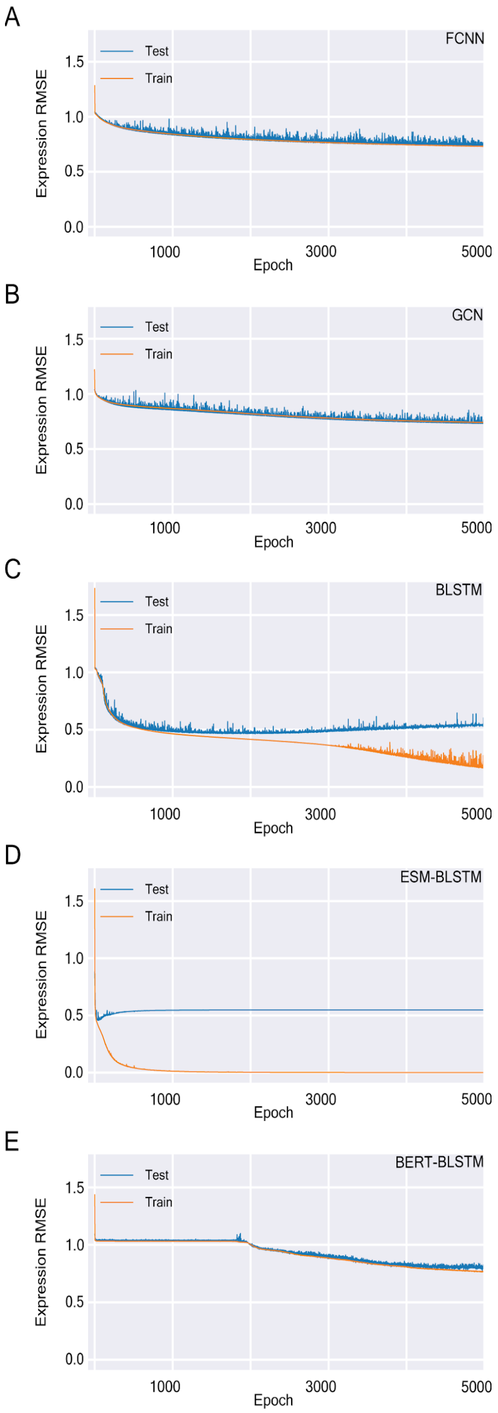

**S Figure 6. Statistics from antibody binding dataset**

A

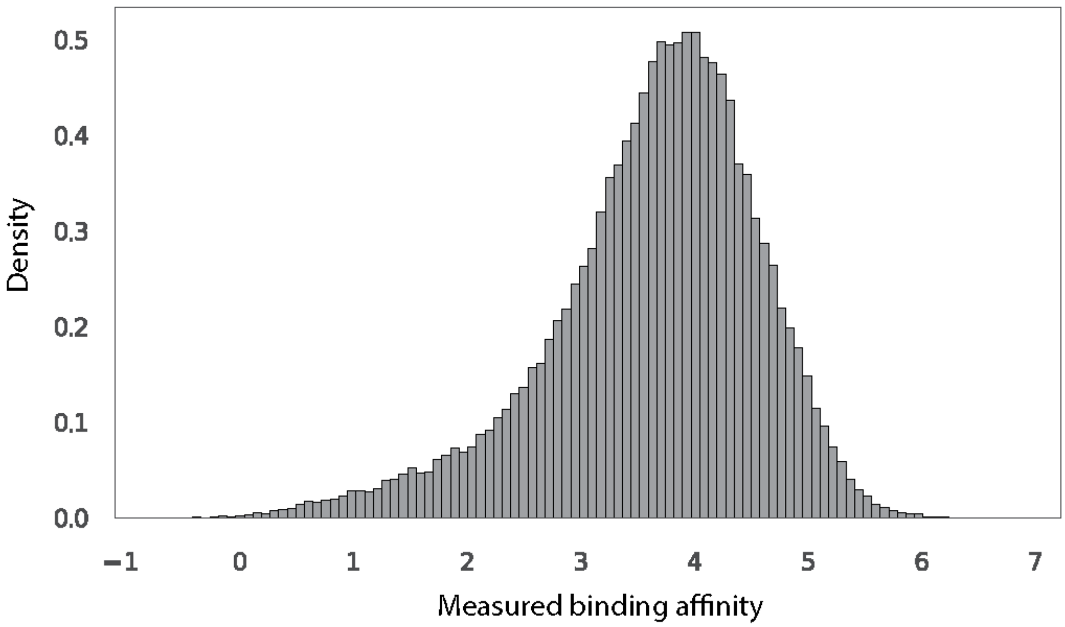


B

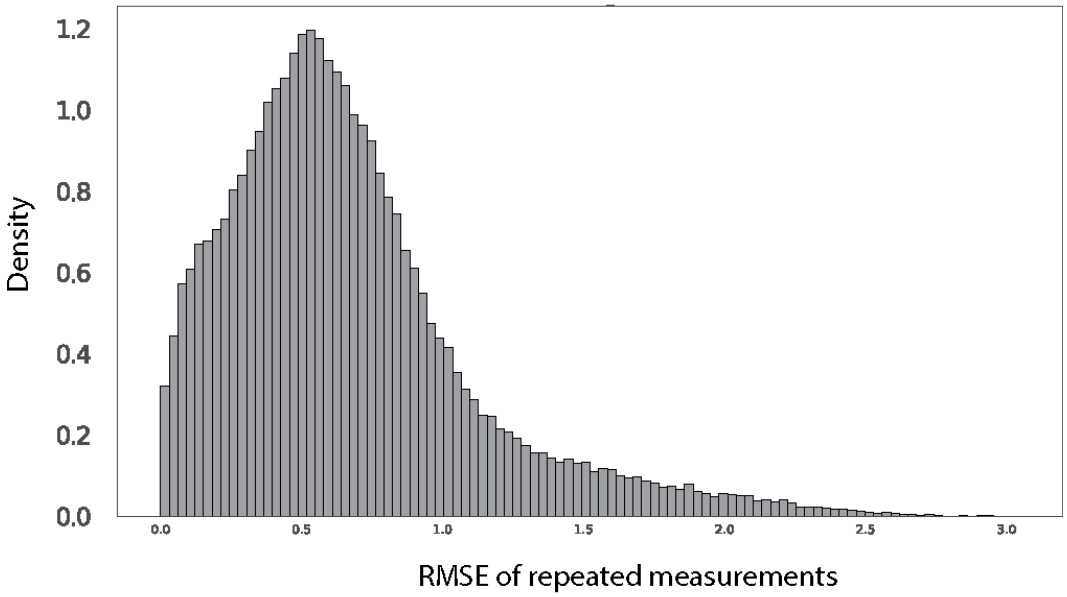

**S Figure 7. Predicting RBD-ACE binding affinities using different models using learned RBD embeddings. A.** Fully connected network with RBD learned embedding. **B.** Graph convolutional network with RBD learned embedding. **C.** Standalone BLSTM model with fixed RBD learned embedding. **D.** BERT-BLSTM model.

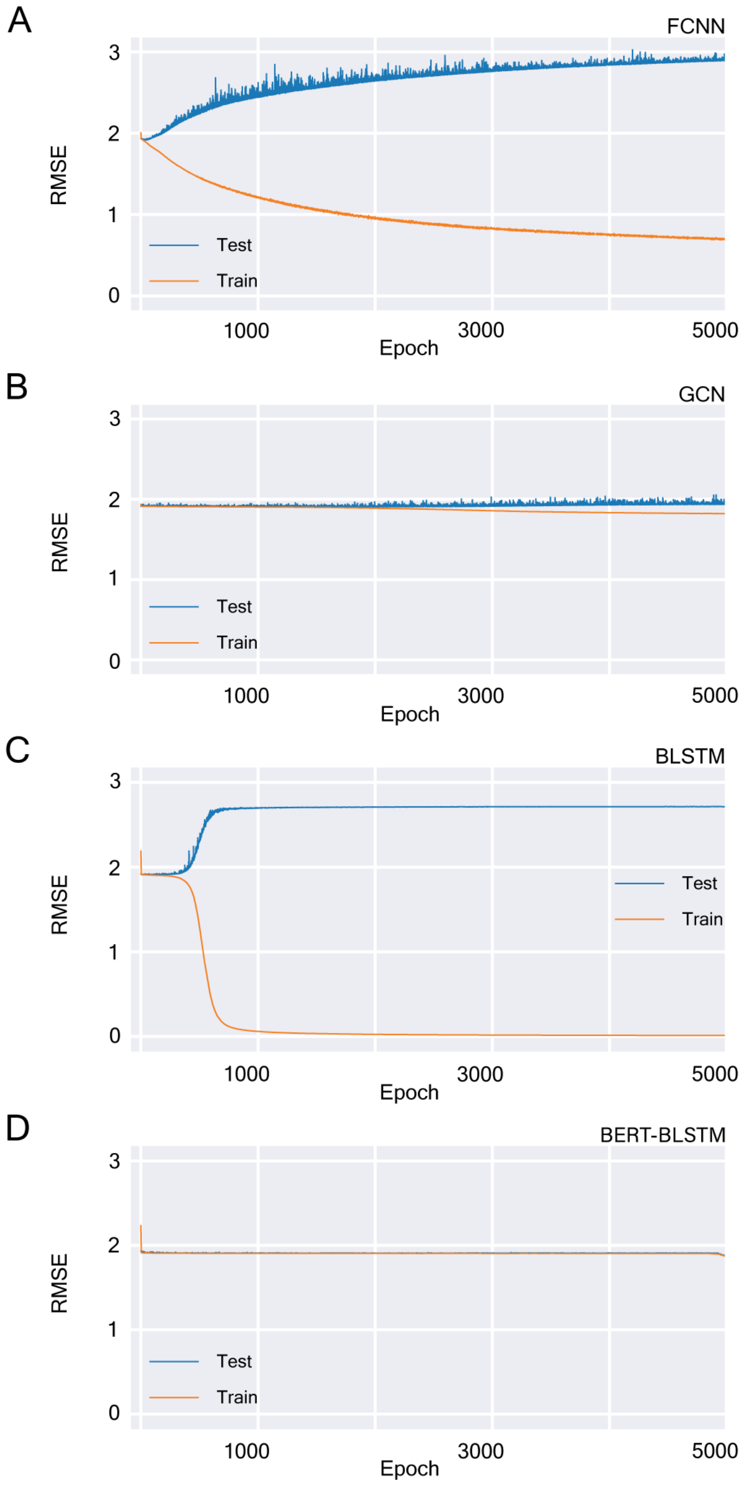

**S Figure 8. Predicting RBD expression levels using different models. A.** Fully connected neural network with RBD learned embedding. **B.** Graph convolution network with RBD learned embedding. **C.** Standalone BLSTM with RBD learned embedding. **D.** BERT-BLSTM model.

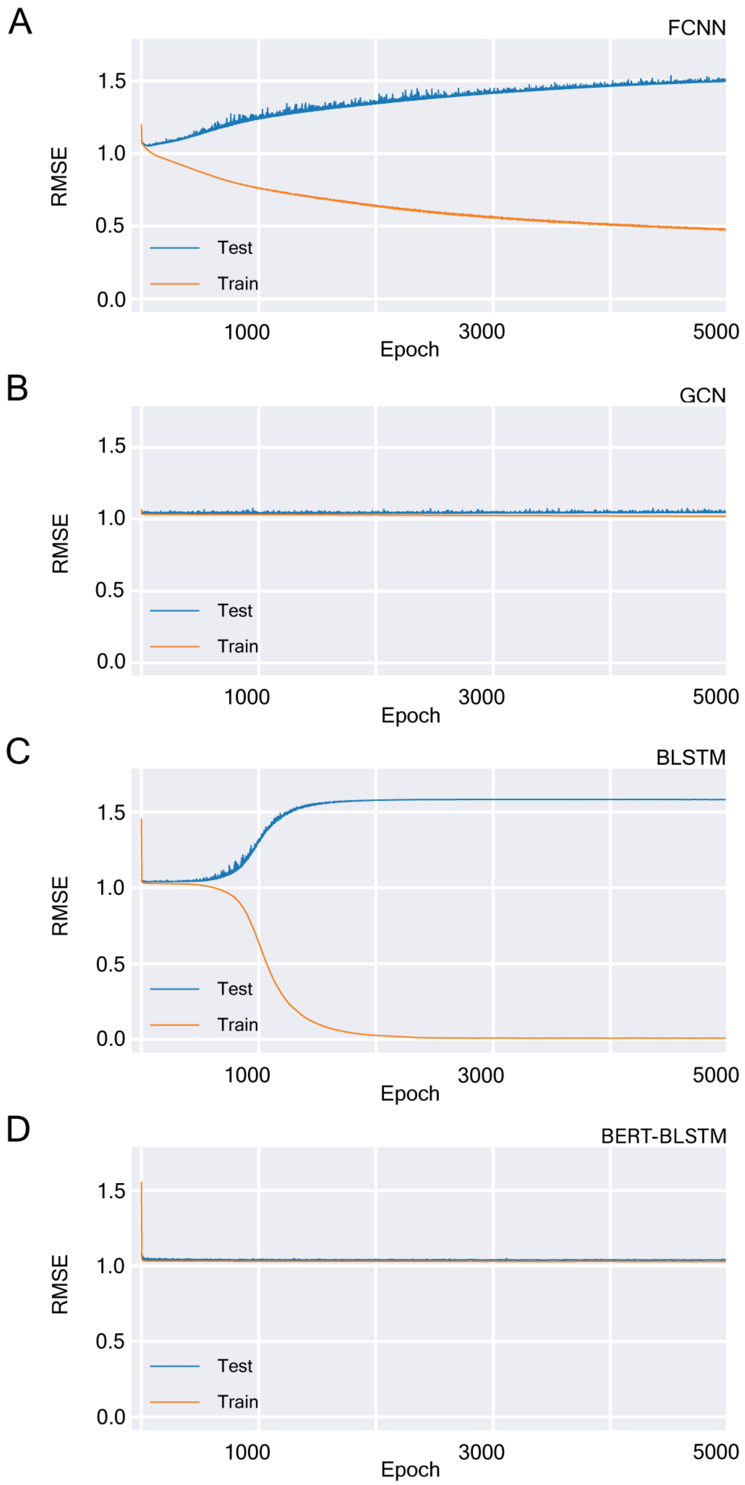

**S Table 1. Lineage composition of three Omicron dominating clusters.** Three clusters from Supplementary Figure 4 were selected with the top 5 Pango lineages extracted. Cluster (%) is the percent of that lineage in a cluster. Lineage (%) is the percent of the sequence of the entire lineage that falls into this cluster.

| Cluster | Top 5 Pango Lineages | Cluster % | Lineage % |
|---|---|---|---|
| 1 | BA.2.12* | 79.885057 | 79.428571 |
| 1 | BA.2 | 4.741379 | 1.731375 |
| 1 | BA.5.2* | 2.011494 | 0.637523 |
| 1 | Unassigned | 1.867816 | 1.444444 |
| 1 | BA.2.11* | 1.293103 | 100.000000 |
| 2 | BA.4.6* | 16.366158 | 80.000000 |
| 2 | BA.5.2* | 15.256588 | 10.018215 |
| 2 | BF.7 | 14.771151 | 81.297710 |
| 2 | BF.7.14* | 5.409154 | 93.975904 |
| 2 | Unassigned | 4.576976 | 7.333333 |
| 3 | BQ.1.1* | 77.922926 | 85.214286 |
| 3 | Unassigned | 3.723057 | 6.333333 |
| 3 | BQ.1.18* | 1.959504 | 90.909091 |
| 3 | BQ.1.25* | 1.502286 | 88.461538 |
| 3 | BA.4.6* | 0.783801 | 4.067797 |
